# Machine learning models identify multimodal measurements highly predictive of transdiagnostic symptom severity for mood, anhedonia, and anxiety

**DOI:** 10.1101/414037

**Authors:** Monika S. Mellem, Yuelu Liu, Humberto Gonzalez, Matthew Kollada, William J. Martin, Parvez Ahammad

**Affiliations:** Data Science Team, BlackThorn Therapeutics, 780 Brannan St., San Francisco, CA, 94103, USA

**Keywords:** Elastic Net, LASSO, Random Forest, Regression, CNS, Depression

## Abstract

*Background:* Insights from neuroimaging-based biomarker research have not yet translated into clinical practice. This translational gap could be because of a focus of psychiatric biomarker research on diagnostic classification, rather than on prediction of transdiagnostic psychiatric symptom severity. Currently, no transdiagnostic, multimodal predictive models of symptom severity that include neurobiological characteristics have been described. *Methods:* We built predictive models of three common symptoms in psychiatric disorders (dysregulated mood, anhedonia, and anxiety) from the Consortium for Neuropsychiatric Phenomics dataset (n=272) which contains clinical scale assessments, resting-state functional-MRI (rs-fMRI) and structural-MRI (sMRI) imaging measures from patients with schizophrenia, bipolar disorder, attention deficit and hyperactivity disorder, and healthy control subjects. We used an efficient, data-driven feature selection approach to identify the most predictive features from these high-dimensional data. *Results:* This approach optimized modeling and explained 65-90% of variance across the three symptom domains, compared to 22% without using the feature selection approach. The top performing multimodal models retained a high level of interpretability which enabled several clinical and scientific insights. First, to our surprise, structural features did not substantially contribute to the predictive strength of these models. Second, the Temperament and Character Inventory scale emerged as a highly important predictor of symptom variation across diagnoses. Third, predictive rs-fMRI connectivity features were widely distributed across many intrinsic resting-state networks (RSN). *Conclusions:* Combining rs-fMRI with select questions from clinical scales enabled high levels of prediction of symptom severity across diagnostically distinct patient groups and revealed that connectivity measures beyond a few intrinsic RSNs may carry relevant information for symptom severity.

## Introduction

The field of psychiatry has long relied on making diagnoses and recommending treatment for disorders based solely on clinical phenomenology, but this approach may hamper prognostic assessment, treatment, and drug development (1, 2). Biomarkers are biological characteristics that can serve as indicators for normal or pathogenic processes or intervention and exposure responses (3). Thus, they enable prediction of these assessments and outcomes and can therefore be an important clinical tool for clinicians. But biomarker development within psychiatry lags behind other areas of medicine. One reason for this may be that the field is still exploring biological measures able to robustly describe a complex psychiatric space. Another reason may be that a diagnostic biomarker approach does not fully account for the heterogeneity of symptoms under the umbrella of a single diagnosis or the shared symptoms between multiple diagnoses. This is because clinical symptoms such as depressed/elevated mood, anhedonia, and anxiety span multiple diagnostic categories (4) but vary between patients, show differential responses to treatments, and follow different prognostic trajectories. As suggested by the Research Domain Criteria (RDoC) approach (5, 6), supplementing the current clinical approach with a biologically-grounded approach that addresses transdiagnostic symptom variation may provide an avenue for creating more robust biomarkers in psychiatry.

Evidence is emerging that biological measures, specifically neuroimaging measures such as electroencephalogram (EEG) and functional magnetic resonance imaging (fMRI), are associated with symptom dimensions that span multiple psychiatric disorders or diverse community samples. Several studies have suggested that derived symptom dimensions are associated with changes in EEG power or resting-state fMRI (rs-fMRI) connectivity transdiagnostically (i.e., where multiple diagnostically-distinct patient groups are modeled together) (7–9). Others have found links between task-based fMRI activation or rs-fMRI connectivity and existing anhedonic, depressive, and anxiety symptom dimensions transdiagnostically (10–13). While the symptom to neurophysiological links in these studies are compelling, most lack a predictive framework, and we are not aware of any attempts toward creating transdiagnostic predictive symptom models integrating whole-brain, circuit-based functional neuroimaging with other data modalities. Combining biological and clinical variables has led to improved predictability in cancer models (14–16) but is underexplored in psychiatry. The predictive framework is especially powerful beyond associative frameworks (such as correlation analyses) as it not only allows multivariate modeling to deal with the high-dimensional, multimodal data but also testing of predictive value and generalizability of those models on a reserved subset or new samples (17–20). Such transdiagnostic, multimodal predictive models, optimized for performance, could eventually be used practically as clinical tools without constraining clinicians by diagnosis.

It is not clear if symptoms have a more circumscribed biological basis to a single or a select few brain networks as proposed in a recent taxonomy (21, 22) or a basis in multiple networks (e.g., (10)), so a whole-brain fMRI connectivity approach could help assess this outstanding question. In particular, we were motivated to test this simultaneously for multiple symptoms to also assess the potential of a short rs-fMRI scan to act as a test for a multi-symptom panel (akin to a blood test). It is also not known if a single, broad self-report clinical assessment (like the Temperament and Character Inventory or the Hopkins Symptom Checklist) or multiple, more specific instruments are better at assessing multiple symptoms. A publicly-available dataset, the Consortium for Neuropsychiatric Phenomics (CNP; (23)) has included 3 patient groups with shared genetic risk (24) and MRI and clinical scale data with which we can assess these questions, so here we used it to explore 3 symptom dimensions – dysregulated mood, anhedonia, and anxiety. But the multimodal, high-dimensional variable space of both MRI and clinical scale data presents both methodological and interpretation challenges in building predictive symptom severity models. Thus, we apply a commonly-used data-driven feature selection approach from the field of machine learning to search through a high-dimensional space and optimize model performance and interpretability. We take an importance-weighted, forward selection approach (a variation of forward-stepwise selection, see (25)) as a data-driven way to identify the optimal feature subset to include in regression model-building (see (26, 27) for similar forward selection subset approaches to fMRI-based modeling). Finding an optimal subset helps in high-dimensional cases where the number of features (usually denoted by *p*) is greater than the number of samples (usually denoted by *n*) to avoid overfitting of the models. It also reduces nuisances from uninformative input variables without requiring the modeler to decide *a priori* whether a variable is signal or noise.

Thus, our main objective in this study was to use a fully data-driven method to find highly-predictive, transdiagnostic multimodal symptom severity models for several measures of dysregulated mood, anhedonia, and anxiety. Critically, demonstration of a set of highly-predictive models for multiple symptoms from a few broader tests (rs-fMRI, sMRI, scales) would suggest a more feasible application of these tests to the clinic than multiple narrower tests (like task-based fMRI). As our approach leads to interpretable features (e.g., the top predictive scale items or rs-fMRI node-to-node connections are known), a secondary objective was to better understand the features returned by the best biomarkers at a category level (scales or intrinsic networks) and compare these to previously proposed hypotheses about associated behavioral and physiological components underlying symptoms. With our multimodal approach, we were also able to evaluate the relative contributions of multiple feature types in the context of multimodal models.

## Methods and Materials

### Participants

Full details on the participants from the publicly-available CNP dataset are available including the consenting and human protections information in the data descriptor publication (23). Briefly, four groups of subjects were included in the sample which was drawn from adults aged 21-50 years: healthy controls (HC, n=130), Schizophrenia patients (SZ, n=50), Bipolar Disorder patient (BD, n=49), and Attention Deficit and Hyperactivity Disorder (ADHD, n=43). Stable medications were permitted for participants. Diagnoses were based on the Structured Clinical Interview for DSM-IV (SCID) and supplemented with the Adult ADHD Interview. Out of all subjects, one had incomplete clinical phenotype data from the clinical scales used in this study, 10 had missing sMRI data, and 10 had missing rs-fMRI data. Fifty-five subjects had a headphone artifact in their sMRI data, whereas 22 subjects had errors in the structural-functional alignment step during MRI preprocessing. These subjects were excluded from the corresponding modeling analyses performed in this study. The participant numbers and demographics information are given in Table 1.

**Table 1.**
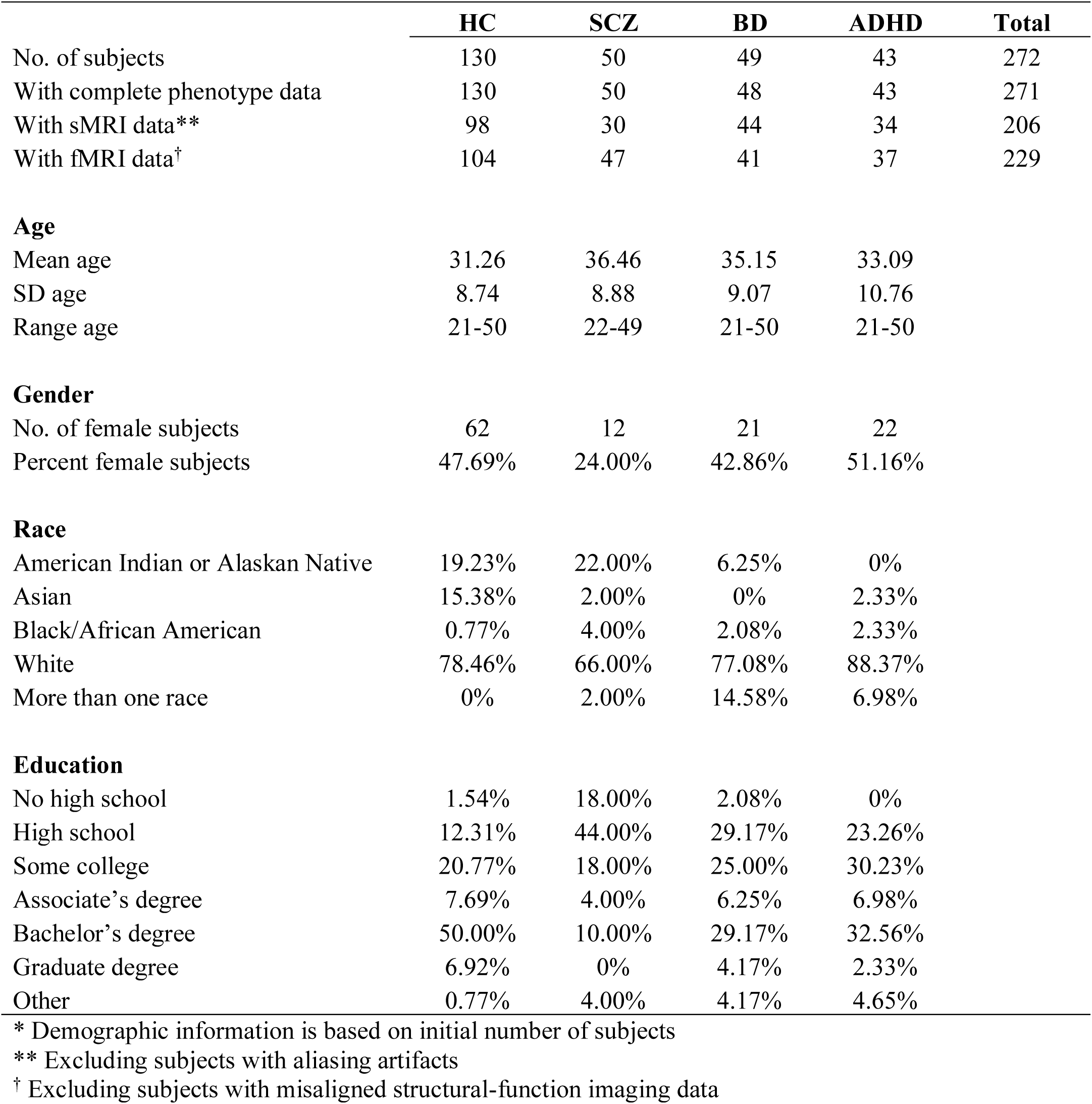
Participant Demographics: Demographics for participants in the CNP dataset broken down by patient group.

### CNP Dataset

The CNP dataset (release 1.0.5) was retrieved from the OpenNeuro platform (https://openneuro.org/datasets/ds000030/versions/00001). Of the extensive behavioral testing that participants underwent, we analyzed results from tests of their self-reported symptoms and traits as clinician-administered instruments were only given to subsets of participants. The self-reported tests used in our analysis include the Chapman social anhedonia scale (denoted *Chapsoc*), Chapman physical anhedonia scale (*Chapphy*), Chapman perceptual aberrations scale (*Chapper*), Chapman hypomanic personality scale, Hopkins symptom checklist (*Hopkins*), temperament and character inventory (*TCI*), adult ADHD self-report scale v1.1 screener (*ASRS*), Barratt impulsiveness scale (*Barratt*), Dickman functional and dysfunctional impulsivity scale (*Dickman*), multidimensional personality questionnaire – control subscale (*MPQ*), Eysenck’s impulsivity inventory (*Eysenck*), scale for traits that increase risk for bipolar II disorder (*Bipolar_ii*), and Golden and Meehl’s Seven MMPI items selected by taxonomic method (*Golden*).

All participants used in this sample also underwent magnetic resonance imaging sessions with T1 scans (structural MRI) and T2* scans of blood-oxygen-level-dependent (BOLD) resting-state functional-MRI and several tasks. Here we only utilize the sMRI and rs-fMRI data (304 seconds in length), and full details of these MRI acquisitions can be found in Poldrack et al. (23). Our decision to focus on rs-fMRI data over task-based fMRI was also due to its ability to provide a fine-grained, data-driven set of functional connectivity features that exhibit meaningful individual differences (28) that relate to symptoms (e.g., (8, 10, 29)).

### Preprocessing Data into Features

We chose to use all responses to individual questions from the 13 self-report scales as input features for a total of 578 questions. Subjects who had missing values for any scales used in a particular model were not included in that model. Outcome variables for modeling mood, anhedonia, and anxiety were also selected from the clinical scales.

Preprocessing of sMRI was performed using the *recon-all* processing pipeline from the Freesurfer software package (30). Briefly, the T1-weighted structural image from each subject was intensity normalized and skull-stripped. The subcortical structures, white matter, and ventricles were segmented and labeled according to the algorithm described in (30). The pial and white matter surfaces were then extracted and tessellated (31), and cortical parcellation was obtained on the surfaces according to a gyral-based anatomical atlas which partitions each hemisphere into 34 regions (32). The structural features from bilateral *aparc.stats* and *aseg.stats* files were extracted via the *aparcstats2table* and *asegstats2table* functions in Freesurfer.

Preprocessing of rs-fMRI was performed using the AFNI software package (33). Preprocessing of each subject’s echo planar image (EPI) data included several steps: removal of the first 3 volumes (before the scanner reached equilibrium magnetization), de-spiking, registration of all volumes to the now first volume, spatial smoothing with a 6 mm full-width half-maximum Gaussian filter, and normalization of all EPI volumes by the mean signal to represent data as percent signal change. Anatomical data also underwent several steps: deobliquing of the T1 data, uniformization of the T1 to remove shading artifacts, skull-stripping of the T1, spatial alignment of the T1 and Freesurfer-segmented and -parceled anatomy to the first volume of the EPI data, and resampling of the Freesurfer anatomy to the resolution of the EPI data. Subsequently, we used the ANATICOR procedure (34) for nuisance tissue regression. White matter and ventricle masks were created and used to extract the BOLD signals (before spatially-smoothing the BOLD signal). A 25mm-radius sphere at each voxel of the white matter mask was used to get averaged local white matter signal estimates while the average ventricle signal was calculated from the whole ventricle mask. Time series for the motion estimates, and the BOLD signals in the ventricles and white matter were detrended with a 4^th^ order polynomial. To clean the BOLD signal, we regressed out the nuisance tissue regressors and the six motion estimate parameters. Cleaned data residuals were used for all subsequent analysis. Both the preprocessed T1 scan and the cleaned residuals of the EPI scan were warped to MNI space and resampled to 2mm isotropic voxels. The time series of the cleaned residual data was extracted from each of 264 regions of interest (ROIs) as delineated by the Power atlas (35). At each ROI, the signals from the voxels within a 5mm radius sphere were averaged. Pearson’s correlations were then calculated between the averaged time series from all ROIs yielding 34716 unique edges in the functional connectivity graph (upper triangle of the full correlation matrix). Quality control (QC) for MRI preprocessing was performed individually on the whole dataset by two authors (MM, YL) who had 85% and 89% agreement between them regarding rejection decisions for each participant’s sMRI and rs-fMRI data, respectively. Specifically, subjects were excluded if they had mis-registration between fMRI and sMRI scans, >3mm head motion in the fMRI scan, headphone artifacts that overlapped with brain tissue in the sMRI scan, incorrect segmentation in the sMRI scan, and aliasing or field of view artifacts in either scan. Discrepancies were resolved between the two authors in order to create a final rejection list of participants.

Input features for each subject came from the three preprocessed datasets: raw scores on the 578 individual items of 13 self-report clinical scales, 270 Freesurfer-calculated structural measurements (including subcortical volume, cortical volume, cortical area, cortical thickness), and 34716 AFNI-calculated functional connectivity scores between individual ROIs. Subsets of these input features were used as predictor variables in subsequent modeling as explained below.

Output variables that were modeled included those which indexed mood, anhedonia, and anxiety. We predicted a mix of total scores and sub-scale sum or average scores from scales that were given to all three patient groups and HCs to retain the largest number of subjects possible in our models. Each of these scores were already calculated and included in the CNP dataset, and we chose to use them rather than determining our own grouping of individual scores as we did not always have access to original scale questions. For mood, we used the average of depression symptom questions 5, 15, 19, 20, 22, 26, 29, 30, 31, 32, and 54 from the Hopkins inventory (precalculated “Hopkins_depression” score, further referenced as *Mood/Dep_Hopkins* in this study) and the sum of mood questions 1-9 from the Bipolar_ii inventory (precalculated “Bipolar_mood” score, further referenced as *Mood_Bipolar* in this study). Anhedonia was derived from total scores on the Chapman Social Anhedonia scale (precalculated “Chapsoc” score, further referenced as *Anhedonia_Chapsoc* in this study) and the Chapman Physical Anhedonia scale (precalculated “Chapphy” score, further referenced as *Anhedonia_Chapphy* in this study). Anxiety was indexed from the sum of Bipolar_ii anxiety questions 24-31 (precalculated “Bipolar_anxiety” score, further referenced as *Anxiety_Bipolar* in this study) and average of anxiety symptom questions 2, 17, 23, 33, 39, and 50 from the Hopkins anxiety score (precalculated “Hopkins_anxiety” score, further referenced as *Anxiety_Hopkins* in this study). Subjects with missing values (“n/a”) for any input or output variables or who did not pass MRI QC were removed from the input set. As different input feature sets were used, different models had different sample sizes. See the samples sizes resulting from this factor in Supplementary Table S1.

### Regression Modeling

All regression modeling was performed with a combination of Python language code and the Python language toolbox scikit-learn (http://scikit-learn.org/stable/index.html). We modeled six different symptom severity scores across the clinical scales. For each of the six models, we used seven combinations of feature types as the inputs to be able to evaluate performance of single and multimodal feature sets. These included clinical scales only, sMRI only, fMRI only, scales+sMRI, scales+fMRI, sMRI+fMRI, and scales+sMRI+fMRI. As input features varied in their mean values and regularized models require normally-distributed data, we scaled each input feature separately to have zero mean and unit variance. We also wanted to explore performance with a variety of modeling algorithms, so for each scale output and feature set input, we used two regularized general linear model regression algorithms – LASSO and Elastic Net – and one non-linear regression model algorithm – Random Forest – for the modeling. These methods have been found to improve prediction accuracy and interpretability over regular regression methods using ordinary least squares. LASSO (36) was the first to use regularization by imposing an L_1_- penalty parameter to force some coefficients to zero; this step introduces model parsimony that benefits interpretability and predictive performance while guarding against overfitting. If predictor variables are correlated, however, the LASSO approach will arbitrarily force only a subset of them to zero which makes interpretation of specific features more difficult. The Elastic Net algorithm (37) uses both L_1_- and L_2_-penalty parameters to better be able to retain groups of correlated predictor variables; this improves interpretability as highly predictive features will not randomly be set to zero thereby diminishing their importance to the model. It is also better suited in cases when the number of predictor variables is much greater than the number of samples (*p*>>*n*). The non-linear regression algorithm Random Forest was also chosen (38) for comparison purposes.

We built 126 (6 outcome variables x 7 predictor variable sets x 3 model algorithms) sets of models (Figure 1a). For each of these sets of models, hyperparameters were tuned using 5-fold cross-validated grid-search on a training set of data (80% of data), and selected hyperparameters were used on a separate evaluation set of data (20% held-out sample). The approaches of cross-validation and of splitting data between training and evaluation data is one way to minimize overfitting in addition with permutation testing which we also performed below (18, 39). The hyperparameter range for LASSO was alpha equal to 0.01, 0.03, and 0.1 (three samples through the log space between 0.01 and 0.1) which is the coefficient of the L_1_ term. Hyperparameter ranges for Elastic Net were alpha equal to 0.01, 0.03, and 0.1, and l1_ratio equal to 0.1, 0.5, and 0.9 which is the mixing parameter used to calculate both L_1_ and L_2_ terms. Hyperparameter ranges for Random Forest included the number of estimators equal to 10 or 100 and the minimum samples at a leaf equal to 1, 5, and 10. The best hyperparameters were chosen from the model that maximized the r^2^ score (coefficient of determination) across the 5-fold cross-validation procedure in the training set and applied to the model of the never-seen evaluation set.

**Figure 1.**
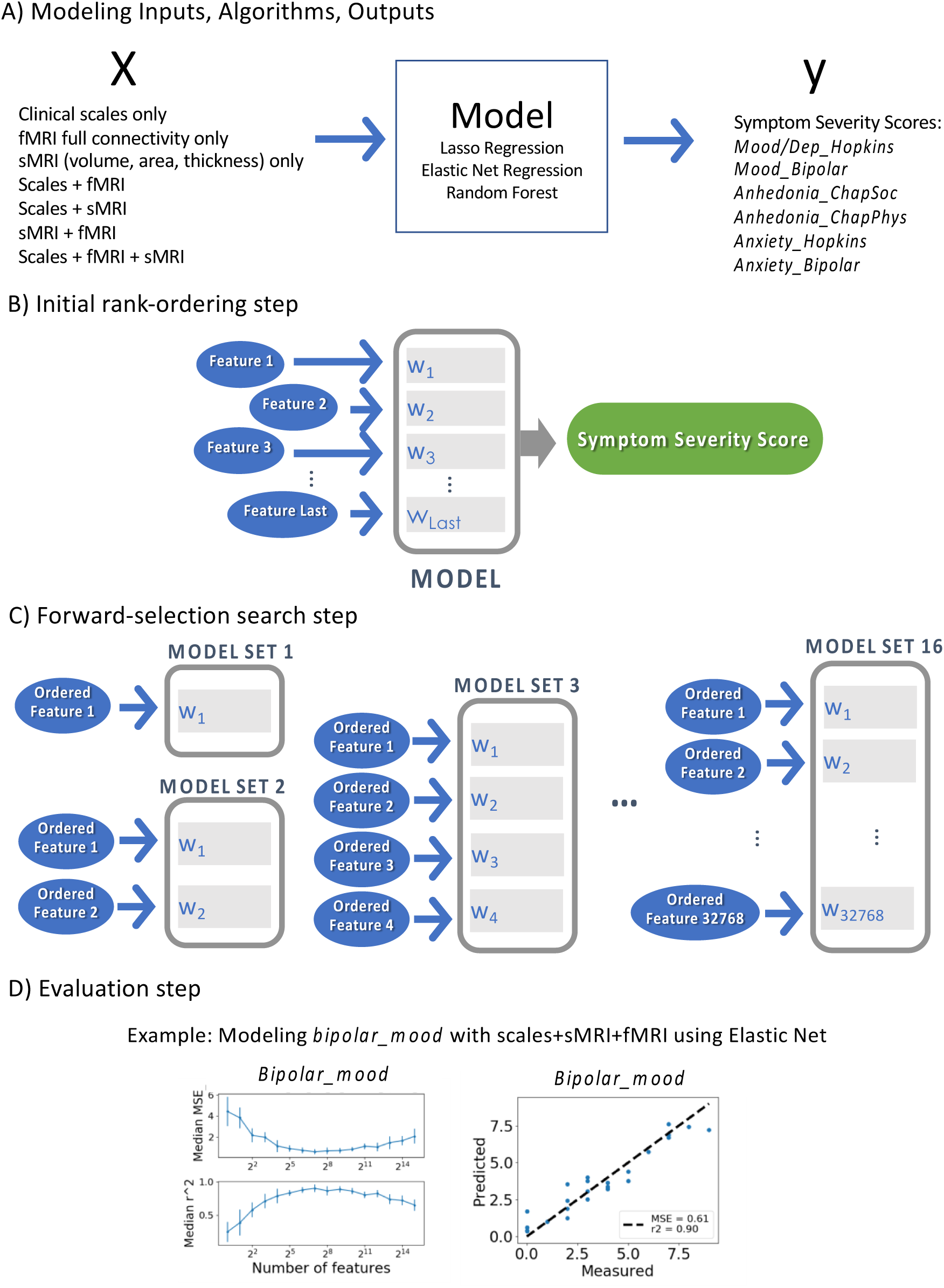
Modeling Approach. a) 126 sets of models were built to examine all permutations of 7 feature set inputs, 3 modeling algorithms, and 6 outcome variables (symptom severity scores). b) The importance-weighted, forward selection approach to regression modeling involved an initial rank-ordering step for ordering features by importance, c) a forward-selection search step for building a series of models utilizing subsets of ordered features selected from the first step, and d) an evaluation step for evaluating each of these models using these candidate subsets according to a prespecified criterion to find the optimal model. An example predicting the total *Mood_Bipolar* score using Elastic Net and scales+sMRI+fMRI as input shows how median MSE (left, top) and median r^2^ (left, bottom) varies with each feature subset, each with standard deviation bars. Measured v. predicted outcome scores (right) show how closely the model predictions are to actual outcome scores for individuals in the held-out sample.

For each of the 126 sets of models, we took an importance-weighted, forward selection approach to regression modeling, involving three main steps: first, an initial rank-ordering step for ordering features by importance; second, a forward-selection search step for building a series of models utilizing growing subsets of ordered features (i.e., the best features) selected from the first step; and third, an evaluation step to choose the best model and subset of features according to a prespecified criterion to find the optimal model (Figure 1b, c, d). This approach thus integrates feature selection into modeling using a multivariate embedded method that can take variable interactions into account to potentially construct more accurate models (40). Within each step, each new model utilized the training/evaluation set split and grid-search procedure to optimize hyperparameters as explained above. First, the feature rank-ordering step uses the full feature set (either scale only, sMRI only, etc.) as the input to the model algorithms which returns not only predicted values for the evaluation dataset but also the importance of each feature for the resulting model (Figure 1b). Feature importance was assessed from the regression coefficients with ordering (most important to least important) based on the absolute value of the coefficient. Ordering by absolute value reflects that features with the largest magnitude influence the symptom severity scores the most. Feature ordering was performed separately for LASSO and Elastic Net models, but as feature importance is harder to assess for the Random Forest algorithm (typical regression coefficients are not available), we used the ordering from the Elastic Net models as input for the subsequent steps of Random Forest modeling instead.

Second, the forward-selection search step systematically searches through subsets of the rank-ordered features (truncated feature sets) for the subset that leads to the best model (Figure 1c). Since having more features than samples (i.e., *p*>>*n*) both increases the risk of overfitting and decreases the performance due to uninformative features adding nuisances, we chose this data-driven way of searching the ordered feature space for an optimal subset of features. We ran a series of regressions on subsets of the ordered features with subsets chosen in powers of 2 (i.e., inputting the top feature only, the top 2 features only, the top 4 features only, etc.) up to 2^15^ features. In order to generate descriptive statistics for this step, we used 25 iterations of modeling for each feature subset to get median and standard deviation metric scores. The metrics chosen for the final step of evaluation were mean squared error (MSE) and r^2^. The median r^2^ and standard deviation of r^2^ were found for each subset. And the “best model” overall was selected by finding the maximum median r^2^ value over all feature subsets and selecting the model that corresponded to that max median r^2^ value (Figure 1d). All subsequent follow up is on the 126 best models for each combination of input x model type x output.

To find which input feature set (clinical scales only, sMRI only, fMRI only, scales+sMRI, scales+fMRI, sMRI+fMRI, and scales+sMRI+fMRI) and which model type (LASSO, Elastic Net, Random Forest) lead to the best biomarkers, subsequent comparisons were also made based on the r^2^ of the best models. The r^2^ is a standardized measurement of explained variance (with a maximum value of 1 but an unbounded minimum) while the MSE values are not standardized across the different models making it less appropriate to use MSE for comparison.

To test alternative hypotheses that modeling may have been impacted by overfitting or variables of no interest, we implemented several control scenarios. We compared model performance for the best models (chosen by the methods above) with models with permuted outcome variables (to test for overfitting) and models that included variables of no interest. In the first case, the null hypothesis is that the features and severity scores are independent; however, an overfit model could misidentify dependence. But if the high performance of our models is due to identification of real structure in the data rather than overfitting, the best models will perform significantly better than models built from the permuted data and we can reject the null hypothesis (41). After the original ordering of features and selection of the 2^n^ subset that led to the best model, we permuted severity scores across subjects for a given outcome variable 100 times and built 100 models based on the permuted scores. We then calculated predictability (assessed with r^2^) from these 100 permuted models which allowed us to generate an empirically-derived distribution of r^2^ values for calculating a test statistic (p-value) compared to the median r^2^ of the chosen best model. In the second control case, models built with only predictor variables of no interest allowed us to assess the predictability of these variables to see if possible confounding variables drive our results. These variables of no interest included age, gender, years of schooling, in-scanner mean framewise displacement which was calculated as an L2 norm, and sharp head motion (output of AFNI’s @1dDiffMag). Modeling was also performed 100 times to generate the r^2^ score distribution and the median r^2^ was compared to this distribution.

## Results

### Selection of models to predict symptom severity

Within a multimodal dataset, we attempted to find the best predictive models for symptom severity transdiagnostically comparing results across models for mood, anhedonia, and anxiety for different predictor variable feature sets and different machine learning algorithms. We found the best r^2^ metric was generally for the scales+sMRI+fMRI input set using Elastic Net across the different outcome variables (a full explanation of how we compared these models for mood, anhedonia and anxiety outcome scores are further explained in the Supplementary Materials). Thus, we have chosen to further examine the features returned for this set of models more closely. Also note that the modeling results for Elastic Net using the full feature sets (not the truncated sets returned by the forward modeling approach) on average explained 22% of the variance while truncated sets explained an average of 78% for the scales+sMRI+fMRI input models (metrics for full features sets are presented in Supplementary Table S10).

For the six models using Elastic Net with scales+sMRI+fMRI input feature set, we evaluated model performance on the held-out test (evaluation) set with measured v. predicted plots (Figure 2) and r^2^ values across models for different outcome variables (Figure 3 and Supplementary Table S9, last column). All 6 models were highly predictive with the variance explained ranging from 65-90%. Next, we compared the proportions of features derived from scale, fMRI, and sMRI feature sets for the best model for each outcome variable both among the whole feature set and the top 25% of features (Figure 4a,b). The best models for *Mood/Dep_Hopkins, Anhedonia_Chapphy*, and *Anxiety_Bipolar* had a roughly equal number of scale and fMRI features while *Anxiety_Hopkins, Anhedonia_Chapsoc*, and *Mood_Bipolar* models had a bias towards fMRI features (Figure 4a). Figure 4b demonstrates though that for many outcome variables there is a disproportionate number of scale features in the top features. There is a paucity of sMRI features in both the these models as only *Anhedonia_Chapphy* had any sMRI features selected by the models. Because of this lack of sMRI features overall, we do not further examine this modality.

**Figure 2.**
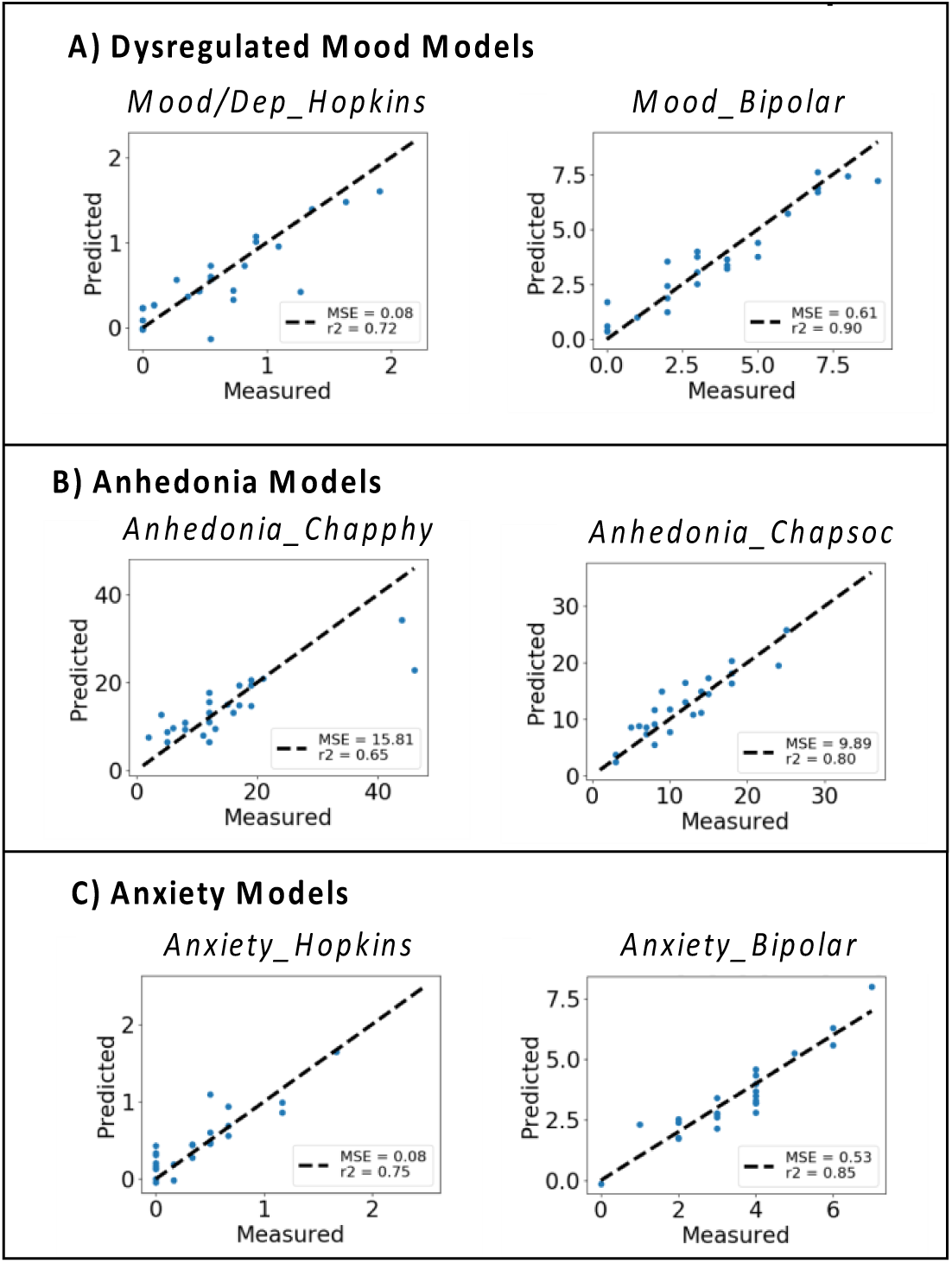
Measured v. predicted values for best models for (A) mood, (B) anhedonia, and (C) anxiety. Each dot in scatter plot represents a single subject from the held-out evaluation set and their measured symptom severity score (x-axis) and predicted score (y-axis). The dashed diagonal line represents a perfect 1-to-1 linear relationship between measured and predicted values. Thus, comparing measured v. predicted outcome scores shows how closely the model predictions are to actual outcome scores for individuals in the held-out samples for this set of models.

**Figure 3.**
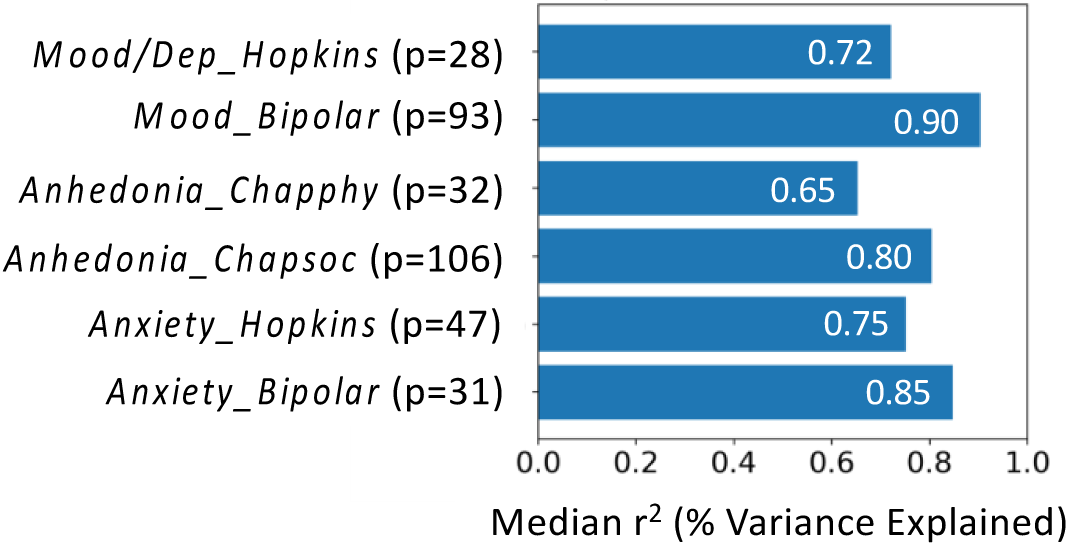
Best median r^2^ for the best models for each outcome variable. Models selected were using Scales+sMRI+fMRI as the input feature set and Elastic Net. Next to each outcome variable, the corresponding number of non-zero features (p) returned by the model appears.

**Figure 4.**
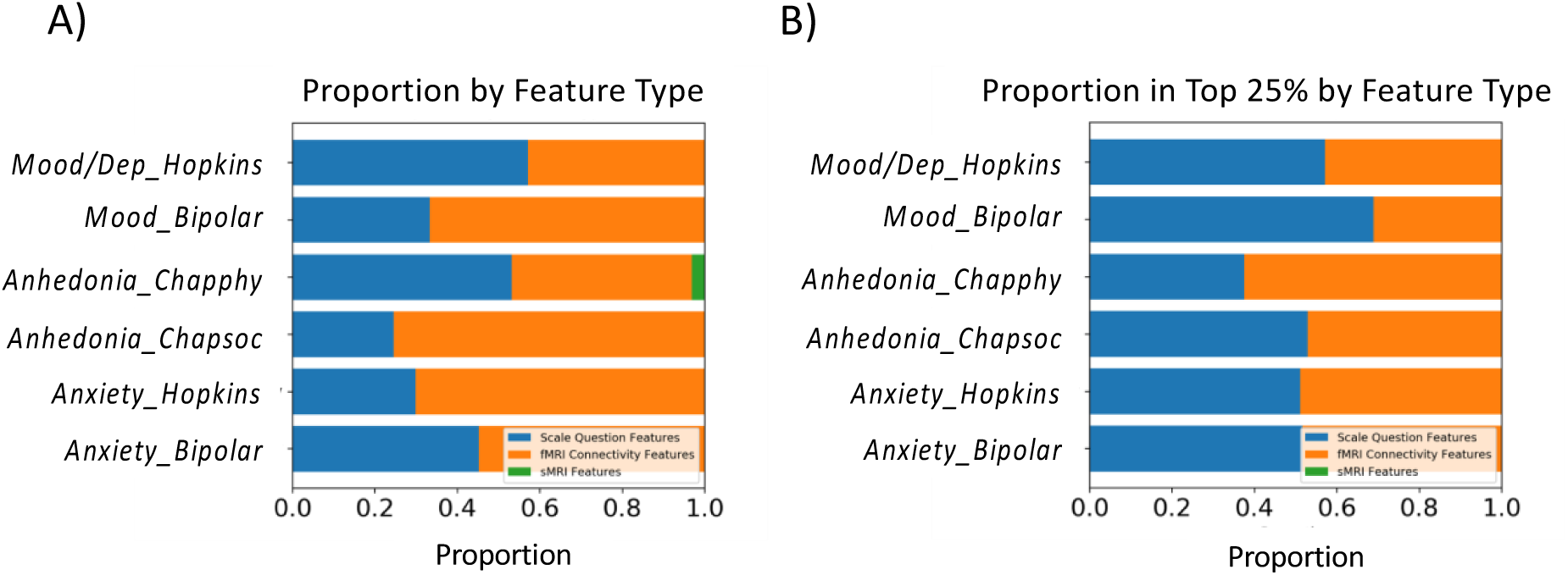
Proportions of feature types in best models. A) Proportion of all features returned by the model. Blue represents proportion of features from scales, orange represents proportion from fMRI connectivity measures, and green represents proportion from sMRI measures. B) Proportion of feature types in the top 25% of features returned by the model showing that most models have equal or greater proportion of scale features than among all the non-zero features.

### Clinical features associated with symptom severity

We can further examine groupings of the scale-based features sorted by proportion of the scales from which they are derived. For each model, the scale features for the best model are proportionately selected from the scales shown in Figure 5. The TCI scale in particular is highly-represented compared to the other scales in all six models (note that Hopkins, Bipolar, Chapphy, and Chapsoc items could not be included in all models as we excluded them when predicting their own sub- or total score). TCI contains a number of questions on temperament and character traits that could be related to a variety of symptoms, and our results suggested that it contains questions that are predictive of mood, anhedonia, and anxiety (Table 2). For example, 43% of the questions predictive of *Anxiety_Bipolar* were from TCI, with the most predictive question being “I am not shy with strangers at all.” Positive responses to this question predicted a lower *Anxiety_Bipolar* score since the regression coefficient was negative in this model. Though not uniformly so, some of the other questions also assessed shyness or worry. The *Anhedonia_Chapsoc* model also had a very high percentage of TCI questions, with the most predictive question being “I would like to have warm and close friends with me most of the time.” Here positive responses indicated decreased social anhedonia severity as the regression coefficient was also negative. While not all questions in the TCI pertain to people and social situations, all but one of the remaining questions that were predictive of the *Anhedonia_Chapsoc* score did include mention of these situations. The predictive questions for the *Anhedonia_Chapphy, Mood/Dep_Hopkins, Mood_Bipolar*, and *Anxiety_Hopkins* scores were more mixed overall though. Full feature lists for the scale items can be requested from the authors. Additionally, Figure 5 shows that *Chaphyp* questions were also predictive in all models (but it only contributed 1-2 items in 5 of the 6 scales). The most numerous questions (6/31) from Chaphyp was for *Mood_Bipolar* which may be expected as the Chapman hypomanic scale and the *Mood_Bipolar* subscore of this scale both include an assessment of mania (as opposed to the *Mood/Dep_Hopkins* score which is more related to depressed mood and depressive symptoms).

**Table 2.**
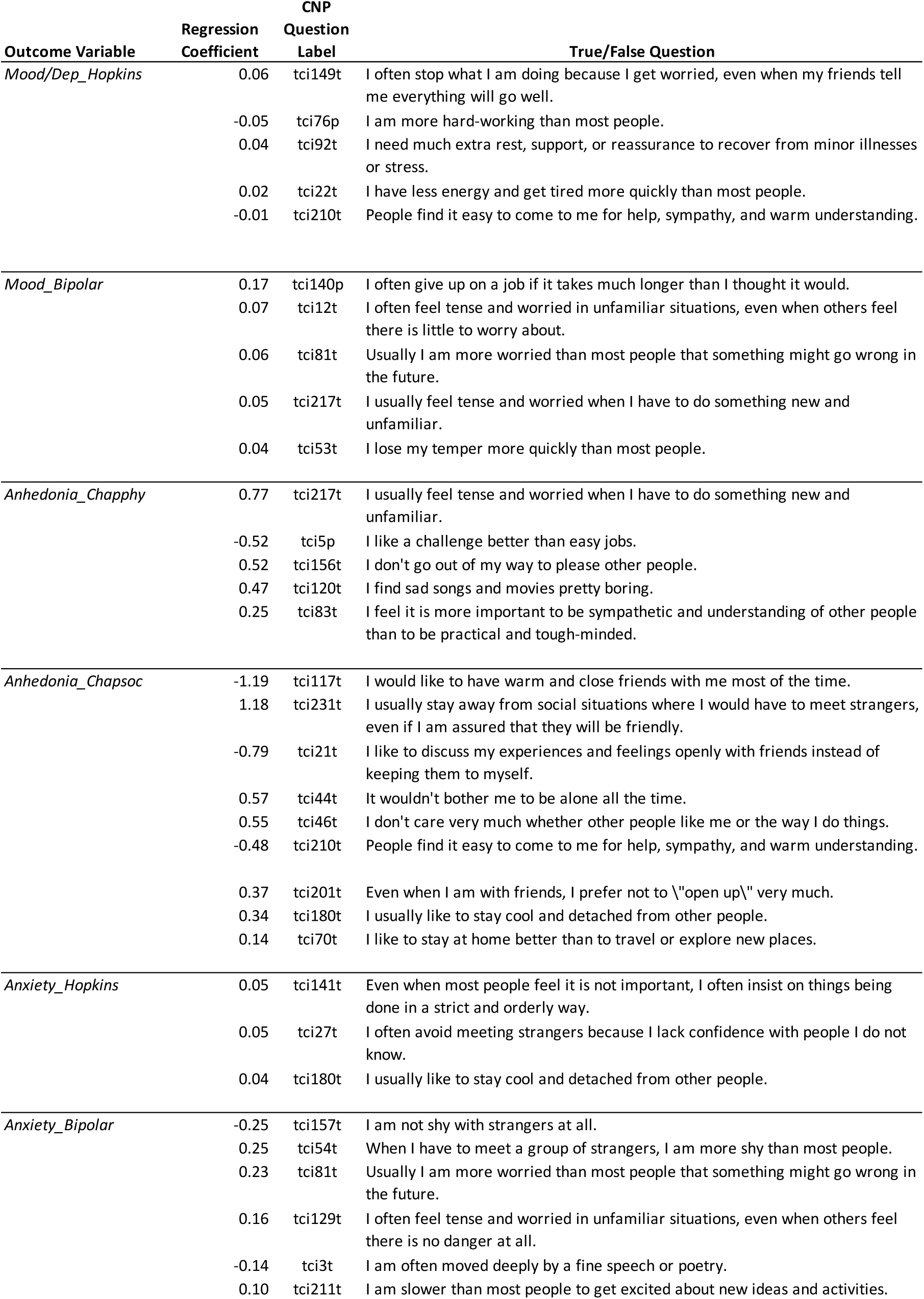
Predictive TCI questions for models of mood, anhedonia, and anxiety. Regression coefficients (ordered by magnitude) are either positive or negative indicating that a “True” answer for the the respective question increased or decreased the outcome variable score, respectively.

**Figure 5.**
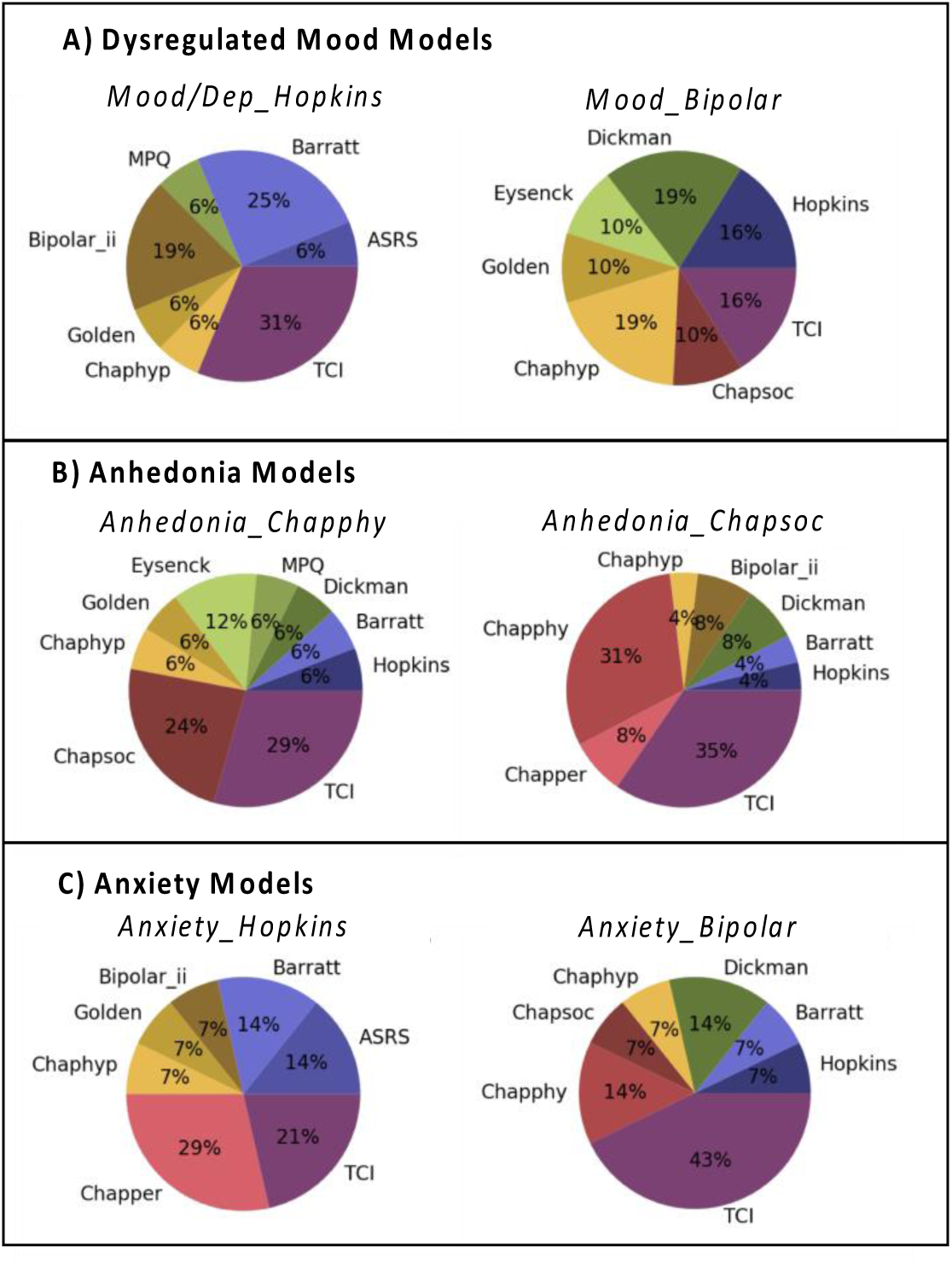
Proportion of features from each scale for the best model predicting (A) mood, (B) anhedonia, and (C) anxiety. Of the features returned by the best model that were scale items, each pie chart shows the proportion of those items that were from the corresponding scales for the model for each outcome variable. For example, for the *Mood/Dep_Hopkins* model, 31% of the scale items were from the *TCI* scale, 6% from the *Chaphyp* scale, etc. Note that this representation of features does not show the sign of the regression coefficient and whether predictive features indicate increasing or decreasing symptom severity.

### Neurobiological characteristics of dysregulated mood, anhedonia and anxiety

The fMRI connectivity features were composed of the strengths of network edges (connections between nodes) but can also be grouped by suggested intrinsic resting-state networks from the Power atlas. As the number of fMRI connectivity features selected by the models were a small subset of all possible fMRI connectivity features, full connectivity matrices are quite sparse (see Supplemental Figure S1). Therefore, we counted the number of edges within and between each intrinsic resting-state network (RSN) and show these counts in the connectivity matrices of Figure 6 (left figure of each panel) for each outcome variable. Please note that connectivity matrices have the same ROIs and networks listed on both axes, and the lower left triangle is redundant to the upper right triangle. The predictive fMRI connectivity features appear mostly distributed across multiple networks rather than selective to a few particular networks as also demonstrated in the color-coded nodes on the brain surfaces (Figure 6, right figure in each panel – please note that cerebellar nodes are not plotted as we only show cortical brain surfaces). Connectivity features implicate nodes in 10 RSNs for *Mood/Dep_Hopkins*, 12 RSNs for *Mood_Bipolar*, 10 RSNs for *Anhedonia_Chapphy*, 12 RSNs for *Anhedonia_Chapsoc*, 13 RSNs for *Anxiety_Hopkins*, and 10 RSNs for *Anxiety_Bipolar* models. Additionally, all 6 models contain features from nodes labeled as “Uncertain” from the Power atlas which indicated that Power et al. were uncertain about the RSN membership of these nodes. While *Anhedonia_Chapsoc* connectivity was also distributed, there was a higher concentration of connectivity features between the Default Mode (DM) network and other networks. In particular, the predictive edges between the DM and other networks mostly originate from the anterior cingulate and/or the medial orbitofrontal lobe. The *Anhedonia_Chapsoc* model also contained nodes in the top 5 features that were located within Williams’ (21) proposed reward circuit including putamen and OFC. Edges either within the DM network or between the DM and other networks consistently were the most numerous features relative to all other within- and between-network features across all models. All the features in each model, including sMRI, are available upon request from the authors.

**Figure 6.**
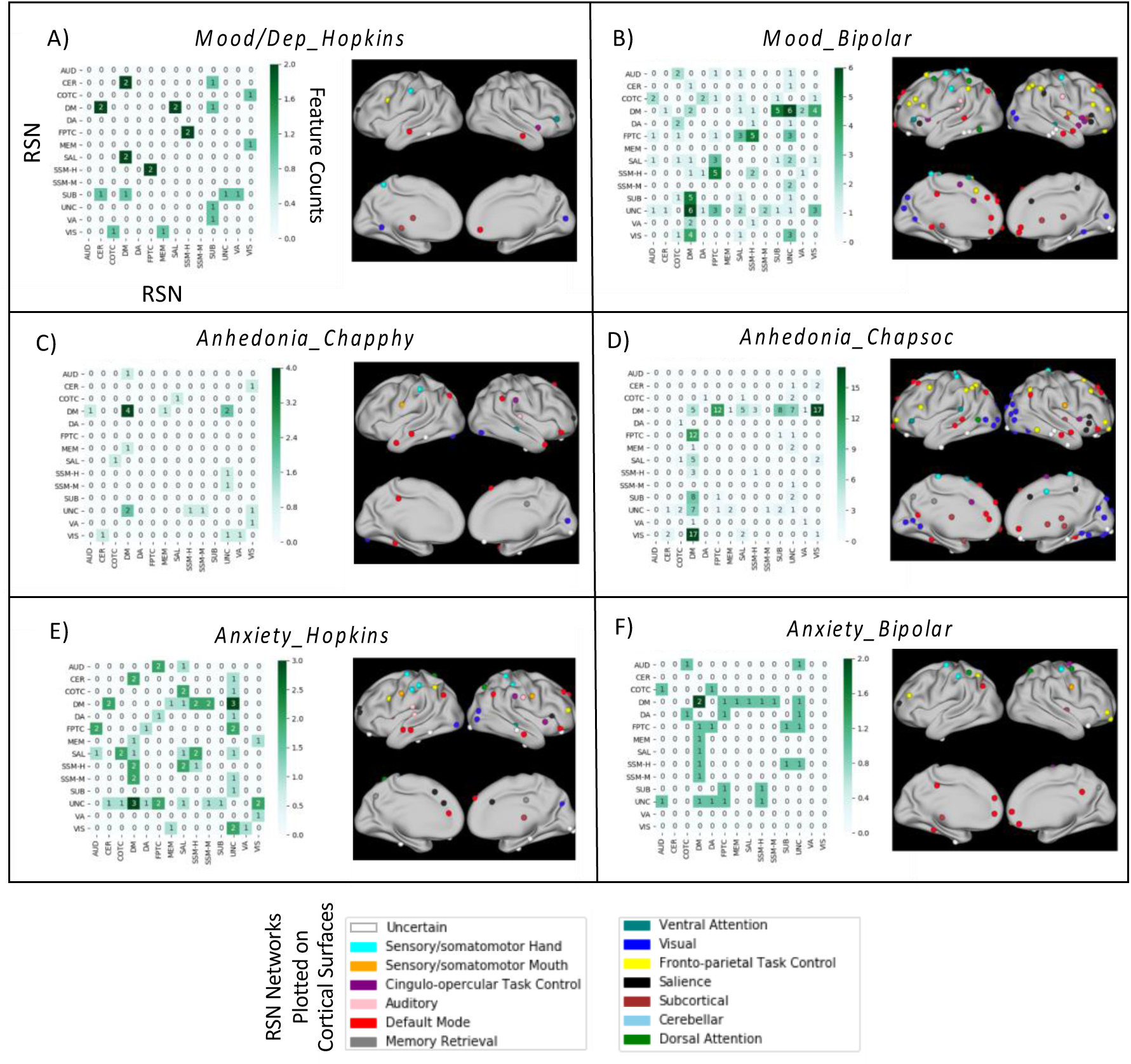
Connectivity Matrices and ROI locations for fMRI connectivity features of best models predicting mood (A and B), anhedonia (C and D), and anxiety (E and F) outcome variables. For all non-zero fMRI connectivity features returned by the respective model, the number of individual edges between two nodes is plotted in the connectivity matrix for that model (left plot in each panel). Each row and column represent a single resting-state network (RSN) from the Power atlas. Thus, darker squares represent more features within or between the given networks with actual feature number superimposed numerically on each square. Upper and lower triangles show redundant information. Cortical surface plots (right plot in each panel) show the ROI locations colored by RSN membership for each model to display the breadth of networks with informative features for each model. Note that since only cortical surfaces are shown, no cerebellar nodes were plotted in the brain plots. Network labels are AUD: Auditory, CER: Cerebellar, COTC: Cingulo-opercular Task Control, DM: Default Mode, DA: Dorsal Attention, FPTC: Fronto-parietal Task Control, MEM: Memory Retrieval, SAL: Salience, SSM-H: Sensory/somatomotor Hand, SSM-M: Sensory/somatomotor Mouth, SUB: Subcortical, UNC: Uncertain (i.e., miscellaneous regions not assigned to a specific RSN), VA: Ventral Attention, VIS: Visual.

### Controls

It is possible that any predictive models generated from the methods above are influenced by confounding demographic or other variables (age, gender, years of schooling, and two measures of in-scanner head motion). For each of the six outcome variables, we built Elastic Net models including these variables of no interest; since the number of predictor variables was only five, the feature selection step was not applied for determining these models though the grid-search, cross-validation, and training/evaluation set steps were applied. In comparison to the median r^2^ values from the best models using scales+sMRI+fMRI features, the predictability of our best models was significantly greater than the variables of no interest models (all p<0.01). Additionally, another set of Elastic Net models with the Scales+sMRI+fMRI feature set but with scrambled severity scores (a permutation testing approach) were built and used to test for overfitting, but we found no evidence of overfitting using this approach as demonstrated by the empirical null distributions (see Supplementary Materials, Figure S2). For all 6 models, the median r^2^ of the best models was statistically significant (p<0.01).

As the models with the least complexity are scales-only models and may be of interest to some readers, we additionally include results for this set of models in the Supplementary Materials section (Supplementary Figure S3 and see Table S9 for metrics of scales-only models built with Elastic Net).

## Discussion

In this study, we explored models for predicting symptom severity of mood disturbances, anhedonia, and anxiety in a transdiagnostic sample. We applied an importance-ranked, forward selection modeling approach to search for the most predictive input features from a set of clinical scale measures, sMRI measures, and rs-fMRI measures. Notably, this data-driven way of selecting feature subsets led to multimodal neurobehavioral models with consistently high predictability across multiple symptom domains and also led to high interpretability since it retains importance scores for individual features. Thus, we demonstrate that the shorter, broadly-applicable 5-minute rs-fMRI scan and a small set of clinical scale assessments can potentially be used to predict a panel of core symptoms commonly found in various psychiatric disorders. Overall, Elastic Net regression models with all three input feature types explained the most variance but features from the different modalities were not equally represented in the models when evaluating the magnitude of the coefficients. The individual, edge-level fMRI connectivity measures between specific network nodes dominated in nearly all of the regression models for different symptom measures, but responses to individual questions in self-report clinical scales were also highly predictive. sMRI measures were not well-represented among the essential features in our models.

As the main objective in this study was to maximize predictability (we chose variance explained as the metric) for models whose regression coefficients could be used for interpretation, we have features that can be assessed for clinical and scientific insights. It must be kept in mind, however, that the features of the best models were chosen in the context of other types of features and are perhaps not as comprehensive as models examining single modalities (i.e., the number of features, *p*, for scale items differs in scales-only models compared to multimodal models). Additionally, the forward selection approach finds that different *p*’s optimize *r*^*2*^ in different models which also restricts comparison across models of similar and different symptom domains. Still, there are potentially useful insights that can be extracted at a high level about each of the feature types and symptom domains.

### Contributions of scale assessments and fMRI connectivity to models

The relative contributions of the different feature types suggest that both scale items and fMRI connectivity were highly important to model predictability. While scale features were not as greatly represented in the best models’ feature list as fMRI features overall, they tended to be more highly represented in the top 25% of features. Thus, their relative importance may be higher than fMRI features, though clearly the multimodal models performed better than scales-only models suggesting that both scale and fMRI components contain unique information. Such a comparison of different feature types in transdiagnostic or community-based symptom severity biomarker studies is not as common (but see for example (42) for a comparison of plasma sterols and demographic variables for prediction of depression severity). It is a valuable step, however, when multiple data types are available for creating predictive models as each data type has benefits and drawbacks in ease of collection, measurement stability, resources required for processing, etc. which may impact choice of data collected for future studies. With regard to the largely negative results for sMRI features, a recent review finds that sMRI regularly underperforms at Major Depressive Disorder (MDD) diagnostic classification in comparison to fMRI (43). Others have also suggested that the lack of studies reporting sMRI abnormalities in SZ, BD, and ADHD has been suggested to reflect the lack of predictability or need for larger sample sizes in detecting effects in this modality (44, 45). It is possible that the sample size in the CNP dataset was not sufficient to maximize the utility of sMRI for prediction.

We further investigated the categorical origins of the fMRI features and clinical scale features for these models. Assessing the categorical groupings of importance-ranked fMRI connectivity features for each model was done according to intrinsic resting-state networks of the Power atlas which partially overlap with a recently proposed taxonomy of symptom-related networks suggested by Williams (21, 22). In contrast with that proposal for a more focused set of brain regions within specific networks as showing impairment that underlie symptoms, this analysis demonstrated that overall these highly-predictive features are distributed across elements of many networks (note that in some models, the edge between two nodes of different intrinsic RSN memberships may be the only element involving those networks) in line with other findings for cross-system representations (46). This may have the implication that it is useful for examining whole-brain connectivity between individual nodes when creating models instead of relying solely upon summary metrics of networks such as graph theory metrics, independent components, or more circumscribed ROI approaches to connectivity (see (10, 47) for a similar full ‘connectome’ approach). Specifically, anhedonia models found not only elements of the reward circuit that Williams (21, 22) proposed as linked to this symptom but also multiple nodes in the DM, Salience (SAL), Cingulo-Opercular Task Control (COTC), Fronto-Parietal Task Control (FPTC), and Visual (VIS) networks among others. Connectivity changes tied to rewarding contexts in this wider set of networks have been observed by others (10, 48) while a meta-analysis of task-based reward processing in MDD demonstrated dysfunctional activation in a broad set of regions including frontal, striatal, cerebellar, visual, and inferior temporal cortex (49). As nodes within the DM network are activated both during self-referential processing and social and emotional processing (29, 50, 51), symptoms that decrease socially pleasurable experiences could have bases in this network. And coordination between several of these networks are necessary for healthy function, but patients with disruptions to the salience network may have trouble switching between DM and executive control networks which may underlie rumination (52) or impaired reward processing (48). Indeed subcortical nodes of the salience network (not included in our analysis) are located in mesocorticolimbic emotional and reward processing centers of the brain (53), so disruption of these functions may propagate to cortical salience regions and beyond.

Our anxiety models also had informative features across a widespread set of networks including high representation in the DM network and sparser representation across executive networks (FPTC, COTC, Dorsal Attention (DA)), SAL network, and sensory networks. Though Williams (21, 22) linked anxiety to a set of networks including a threat circuit, the SAL, DM, and Attention networks, our findings seem to point to a broader set that have some support from previous studies. For example, anxiety has been found related to dysfunction in the DM, SAL, and Somatomotor networks (54), the FPTC network (55), in addition to COTC and Visual Attention (VA) networks (56). Given the core processes of these networks, the underlying elements of anxiety – trouble regulating emotion in fearful situations, detecting and controlling conflict, increased attention to emotional stimuli – have reasonable relationships to this set of networks.

Our depression and mood models predicted outcome variables that were perhaps not as narrowly-focused on a single symptom. The *Mood/Dep_Hopkins* subscore contained depressed mood questions but also ones about guilt, suicide, loss of interest, and somatic concerns, while *Mood_Bipolar* contained questions about both depressed and manic moods, states which the brain may reflect differently (57). We found that both models also relied on a broad set of networks beyond the negative affective circuit (ACC, mPFC, insula, and amygdala) proposed by Williams (21, 22). Both anterior and posterior nodes of the DM network were informative to the model as well as FPTC, COTC, Attention, SAL, and Sensory networks. Cognitive Control networks, Salience, and Attention, and Affective networks have been proposed to be involved in depressed mood (58, 59) while a central node, the subgenual cingulate, is involved in mood (60, 61) and connected within the DM network (62) and approximately observed in our findings. Spielberg and colleagues (63) recently examined connectivity during both depressed and elevated mood in BD and found increased amygdala-sensory connectivity and abnormal prefrontal-parietal connectivity during manic states and extensive orbito-frontal to subcortical and cortical connectivity in depressed states, while Martino and colleagues (64) found the ratio of DM to sensory-motor network activity was greater in a depressed state of BD and less in manic states in BD. Thus a wide set of regions and networks may be involved in depressed and elevated mood, but there may be some dissociation between the two with more DM in depressed mood and sensory involvement in elevated mood. The *Mood_Bipolar* outcome variable included both, so this model includes multiple nodes from both sets of RSNs as important features.

The categorical origins of the clinical scale features demonstrated that there was also some similarity in the scales from which models for the six outcome variables were drawn as most included items from the TCI, Hopkins Symptom Checklist, and the Chapman scales. The TCI in particular was consistently one of the most predictive scales as assessed by number of questions contributed for all six models of mood, anhedonia, and anxiety. This scale measures temperaments such as harm avoidance and novelty seeking (65) which have previously been associated with depression (e.g., (66) and anxiety (67)). Our results suggest that it also contains items predictive of anhedonia. In particular, our models picked out questions from TCI that pertained to social situations as predictive of the social anhedonia severity, but the correspondence between physical anhedonia and the predictive TCI questions was less clear. The consistent representation of TCI across the six measures of three symptoms might suggest that it could be a useful self-report questionnaire to include when screening patients for multiple symptom domains.

### Limitations

Our study has some limitations. Outside of a training and evaluation split of the data, we were not able to perform additional validation of the models’ predictive ability on an external dataset. This would be difficult since the CNP dataset has extensive behavioral phenotyping with clinical scales which few datasets could replicate unless collected prospectively. As we used many of these scales in our models, a thorough validation of the best models would require a dataset which has the same scales. Also, as our models are linear, they only model and select as informative the features which trend in the same direction for all subjects. For example, it has been noted by Whitton et al. (68) that the direction of reward-related activity in the ventral striatum can differ between patient groups that both have anhedonia (i.e., lower in MDD and higher in BD compared to HC). Linear models would not be able to pick out this feature. Thus, these modeling choices constrain our interpretations, and the features that our models return should not be seen as a completely comprehensive list of all informative features for predicting symptom severity or as the only ones potentially involved in the underlying biological processes.

### Conclusions

While we have developed models that identify potential biomarkers for depressed/elevated mood, anhedonia, and anxiety, our work is still highly exploratory. Robust biomarkers must undergo several stages of development requiring replication across differing conditions (e.g., MRI scanners and sites) and increasing sample sizes of datasets to achieve population-level utility (69). This study was able to demonstrate one possible data-driven way to improve biomarker development for predicting symptom severity transdiagnostically and potentially moves us closer to a personalized medicine approach in diagnosing and treating behavioral disorders. Taking a transdiagnostic symptom-based approach may ultimately provide more options for predicting longitudinal and treatment outcomes beyond those afforded by diagnosis alone. Such an approach can possibly loosen the constraints of diagnoses by allowing clinicians to estimate symptom severity in broader populations without a diagnosis. Still, the RDoC framework suggests that “the critical test is how well the new molecular and neurobiological parameters predict prognosis or treatment response” (6), and while the high performance of our symptom severity biomarkers is a step towards creating highly-predictive models for potentially unknown symptom measures, a critical next step would be to create longitudinal ones that predict future measures.

## Acknowledgments

This work was supported by BlackThorn Therapeutics and has been placed on the preprint server BioRxiv. The authors would like to thank Ariana Anderson for early discussions and Clark Gao, Annette Madrid, Atul Mahableshwarkar, Lori Jean Van Orden, Simone Krupka, and Alan Anticevic for discussions and comments on earlier drafts of the manuscript.

## References

1. Joyce DW, Kehagia AA, Tracy DK, Proctor J, Shergill SS (2017): Realising stratified psychiatry using multidimensional signatures and trajectories. J Transl Med. 15: 15.

2. Insel TR, Cuthbert BN (2015): Brain disorders? Precisely. Science. 348: 499–500.

3. (2018, May 2): BEST (Biomarkers, EndpointS, and other Tools) Resource - NCBI Bookshelf. Retrieved from https://www.ncbi.nlm.nih.gov/books/NBK326791/.

4. Kring AM (2008): Emotion disturbances as transdiagnostic processes in psychopathology. Handbook of emotion. 3.

5. Abi-Dargham A, Horga G (2016): The search for imaging biomarkers in psychiatric disorders. Nature Medicine. 22: 1248–1255.

6. Insel T, Cuthbert B, Garvey M, Heinssen R, Pine DS, Quinn K, et al. (2010): Research Domain Criteria (RDoC): Toward a New Classification Framework for Research on Mental Disorders. Am J Psychiat. 167: 748–751.

7. Grisanzio KA, Goldstein-Piekarski AN, Wang M, Ahmed AP, Samara Z, Williams LM (2017): Transdiagnostic Symptom Clusters and Associations With Brain, Behavior, and Daily Function in Mood, Anxiety, and Trauma Disorders. Jama Psychiatry. doi:10.1001/jamapsychiatry.2017.3951.

8. Xia C, Ma Z, Ciric R, Gu S, Betzel RF, Kaczkurkin AN, et al. (2018): Linked dimensions of psychopathology and connectivity in functional brain networks. Nat Commun. 9: 3003.

9. Elliott ML, Romer A, Knodt AR, Hariri AR (2018): A Connectome Wide Functional Signature of Transdiagnostic Risk for Mental Illness. Biol Psychiat. doi:10.1016/j.biopsych.2018.03.012.

10. Sharma A, Wolf DH, Ciric R, Kable JW, Moore TM, Vandekar SN, et al. (2017): Common Dimensional Reward Deficits Across Mood and Psychotic Disorders: A Connectome-Wide Association Study. Am J Psychiat. 174: 657–666.

11. Hägele C, Schlagenhauf F, Rapp M, Sterzer P, Beck A, Bermpohl F, et al. (2015): Dimensional psychiatry: reward dysfunction and depressive mood across psychiatric disorders. Psychopharmacology. 232: 331–341.

12. Satterthwaite T, Cook P, Bruce S, Conway C, Mikkelsen E, Satchell E, et al. (2015): Dimensional depression severity in women with major depression and post-traumatic stress disorder correlates with fronto-amygdalar hypoconnectivty. Mol Psychiatr. 21: 894–902.

13. Yang Z, Gu S, Honnorat N, Linn KA, Shinohara RT, Aselcioglu I, et al. (2018): Network changes associated with transdiagnostic depressive symptom improvement following cognitive behavioral therapy in MDD and PTSD. Mol Psychiatr. 1–10.

14. Pittman J, Huang E, Dressman H, Horng C-F, Cheng SH, Tsou M-H, et al. (2004): Integrated modeling of clinical and gene expression information for personalized prediction of disease outcomes. P Natl Acad Sci Usa. 101: 8431–8436.

15. Nevins JR, Huang ES, Dressman H, Pittman J, Huang AT, West M (2003): Towards integrated clinico-genomic models for personalized medicine: combining gene expression signatures and clinical factors in breast cancer outcomes prediction. Hum Mol Genet. 12: R153–R157.

16. Beane J, Sebastiani P, Whitfield TH, Steiling K, Dumas Y-M, Lenburg ME, Spira A (2008): A Prediction Model for Lung Cancer Diagnosis that Integrates Genomic and Clinical Features. Cancer Prev Res. 1: 56–64.

17. Dubois J, Adolphs R (2016): Building a Science of Individual Differences from fMRI. Trends Cogn Sci. 20: 425–443.

18. Yarkoni T, Westfall J (2017): Choosing Prediction Over Explanation in Psychology: Lessons From Machine Learning. Perspect Psychol Sci. 12: 1100–1122.

19. Lo A, Chernoff H, Zheng T, Lo S-H (2015): Why significant variables aren’t automatically good predictors. Proc National Acad Sci. 112: 13892–13897.

20. Bzdok D, Meyer-Lindenberg A (2017): Machine Learning for Precision Psychiatry: Opportunities and Challenges. Biological Psychiatry Cognitive Neurosci Neuroimaging. doi:10.1016/j.bpsc.2017.11.007.

21. Williams LM (2017): Defining biotypes for depression and anxiety based on large scale circuit dysfunction: a theoretical review of the evidence and future directions for clinical translation. Depress Anxiety. 34: 9–24.

22. Williams LM (2016): Precision psychiatry: a neural circuit taxonomy for depression and anxiety. Lancet Psychiatry. 3: 472–480.

23. Poldrack RA, Congdon E, Triplett W, Gorgolewski KJ, Karlsgodt KH, Mumford JA, et al. (2016): A phenome-wide examination of neural and cognitive function. Sci Data. 3: 160110.

24. Consortium T, Anttila V, Bulik-Sullivan B, Finucane HK, Walters RK, Bras J, et al. (2018): Analysis of shared heritability in common disorders of the brain. Science. 360: eaap8757.

25. Hastie T, Tibshirani R, Friedman J (2009): The elements of statistical learning: data mining, inference, and prediction, 2nd ed. Springer.

26. Gheiratmand M, Rish I, Cecchi GA, Brown MR, Greiner R, Polosecki PI, et al. (2017): Learning stable and predictive network-based patterns of schizophrenia and its clinical symptoms. Npj Schizophrenia. 3: 22.

27. Osuch E, Gao S, Wammes M, Théberge J, Willimason P, Neufeld R, et al. (2018): Complexity in mood disorder diagnosis: fMRI connectivity networks predicted medication-class of response in complex patients. Acta Psychiat Scand. doi:10.1111/acps.12945.

28. Biswal BB, Mennes M, Zuo X-N, Gohel S, Kelly C, Smith SM, et al. (2010): Toward discovery science of human brain function. Proc Natl Acad Sci. 107: 4734–4739.

29. Gotts SJ, Simmons KW, Milbury LA, Wallace GL, Cox RW, Martin A (2012): Fractionation of social brain circuits in autism spectrum disorders. Brain. 135: 2711–2725.

30. Fischl B, Salat DH, Busa E, Albert M, Dieterich M, Haselgrove C, et al. (2002): Whole Brain Segmentation Automated Labeling of Neuroanatomical Structures in the Human Brain. Neuron. 33: 341–355.

31. Fischl B, Dale AM (2000): Measuring the thickness of the human cerebral cortex from magnetic resonance images. Proc National Acad Sci. 97: 11050–11055.

32. Desikan RS, Ségonne F, Fischl B, Quinn BT, Dickerson BC, Blacker D, et al. (2006): An automated labeling system for subdividing the human cerebral cortex on MRI scans into gyral based regions of interest. Neuroimage. 31: 968–980.

33. Cox RW (1996): AFNI: Software for Analysis and Visualization of Functional Magnetic Resonance Neuroimages. Comput Biomed Res. 29: 162–173.

34. Jo H, Saad ZS, Simmons KW, Milbury LA, Cox RW (2010): Mapping sources of correlation in resting state FMRI, with artifact detection and removal. Neuroimage. 52: 571–582.

35. Power JD, Cohen AL, Nelson SM, Wig GS, Barnes K, Church JA, et al. (2011): Functional Network Organization of the Human Brain. Neuron. 72: 665–678.

36. Tibshirani R (1996): Regression shrinkage and selection via the lasso. Journal of the Royal Statistical Society, Series B. 58: 267–288.

37. Zou H, Hastie T (2005): Regularization and variable selection via the elastic net. J Royal Statistical Soc Ser B Statistical Methodol. 67: 301–320.

38. Breiman L (2001): Random Forests. Mach Learn. 45: 5–32.

39. Reddan MC, Lindquist MA, Wager TD (2017): Effect Size Estimation in Neuroimaging. Jama Psychiatry. doi:10.1001/jamapsychiatry.2016.3356.

40. Saeys Y, Inza I, Larrañaga P (2007): A review of feature selection techniques in bioinformatics. Bioinformatics. 23: 2507–2517.

41. Ojala M, Garriga GC (2010): Permutation Tests for Studying Classifier Performance. Journal of Machine Learning Research. 11: 1833–1863.

42. Cenik B, Cenik C, Snyder MP, Brown SE (2017): Plasma sterols and depressive symptom severity in a population-based cohort. Plos One. 12: e0184382.

43. Gao S, Calhoun VD, Sui J (2018): Machine learning in major depression: From classification to treatment outcome prediction. Cns Neurosci Ther. doi:10.1111/cns.13048.

44. Mateos-Pérez J, Dadar M, Lacalle-Aurioles M, Iturria-Medina Y, Zeighami Y, Evans AC (2018): Structural neuroimaging as clinical predictor: A review of machine learning applications. Neuroimage Clin. doi:10.1016/j.nicl.2018.08.019.

45. Goodkind M, Eickhoff SB, Oathes DJ, Jiang Y, Chang A, Jones-Hagata LB, et al. (2015): Identification of a Common Neurobiological Substrate for Mental Illness. Jama Psychiatry. 72: 305–315.

46. Chang LJ, Gianaros PJ, Manuck SB, Krishnan A, Wager TD (2015): A Sensitive and Specific Neural Signature for Picture-Induced Negative Affect. Plos Biol. 13: e1002180.

47. Shen X, Finn ES, Scheinost D, Rosenberg MD, Chun MM, Papademetris X, Constable TR (2017): Using connectome-based predictive modeling to predict individual behavior from brain connectivity. Nat Protoc. 12: 506–518.

48. Yang Y, Zhong N, Imamura K, Lu S, Li M, Zhou H, et al. (2016): Task and Resting-State fMRI Reveal Altered Salience Responses to Positive Stimuli in Patients with Major Depressive Disorder. Plos One. 11: e0155092.

49. Zhang W-N, Chang S-H, Guo L-Y, Zhang K-L, Wang J (2013): The neural correlates of reward-related processing in major depressive disorder: A meta-analysis of functional magnetic resonance imaging studies. J Affect Disorders. 151: 531–539.

50. Li W, Mai X, Liu C (2014): The default mode network and social understanding of others: what do brain connectivity studies tell us. Front Hum Neurosci. 8: 74.

51. Mars RB, Neubert F-X, Noonan MP, Sallet J, Toni I, Rushworth MF (2012): On the relationship between the “default mode network” and the “social brain.” Front Hum Neurosci. 6: 189.

52. Belleau EL, Taubitz LE, Larson CL (2015): Imbalance of default mode and regulatory networks during externally focused processing in depression. Soc Cogn Affect Neur. 10: 744–751.

53. Menon V (2015): Brain Mapping. Syst. 597–611.

54. Peterson A, Thome J, Frewen P, Lanius RA (2013): Resting-State Neuroimaging Studies: A New Way of Identifying Differences and Similarities among the Anxiety Disorders? Can J Psychiatry. 59: 294–300.

55. Cole MW, Repovš G, Anticevic A (2014): The Frontoparietal Control System. Neurosci. 20: 652–664.

56. Sylvester CM, Corbetta M, Raichle ME, Rodebaugh TL, Schlaggar BL, Sheline YI, et al. (2012): Functional network dysfunction in anxiety and anxiety disorders. Trends Neurosci. 35: 527–535.

57. Chase HW, Phillips ML (2016): Elucidating Neural Network Functional Connectivity Abnormalities in Bipolar Disorder: Toward a Harmonized Methodological Approach. Biological Psychiatry Cognitive Neurosci Neuroimaging. 1: 288–298.

58. Fischer AS, Keller CJ, Etkin A (2016): The Clinical Applicability of Functional Connectivity in Depression: Pathways Toward More Targeted Intervention. Biological Psychiatry Cognitive Neurosci Neuroimaging. 1: 262–270.

59. Kaiser RH, Andrews-Hanna JR, Wager TD, Pizzagalli DA (2015): Large-Scale Network Dysfunction in Major Depressive Disorder: A Meta-analysis of Resting-State Functional Connectivity. Jama Psychiatry. 72: 603–611.

60. Mayberg HS, Lozano AM, Voon V, McNeely HE, Seminowicz D, Hamani C, et al. (2005): Deep Brain Stimulation for Treatment-Resistant Depression. Neuron. 45: 651–660.

61. Mayberg HS, Liotti M, Brannan SK, McGinnis S, Mahurin RK, Jerabek PA, et al. (1999): Reciprocal Limbic-Cortical Function and Negative Mood: Converging PET Findings in Depression and Normal Sadness. Am J Psychiat. 156: 675–682.

62. Greicius MD, Flores BH, Menon V, Glover GH, Solvason HB, Kenna H, et al. (2007): Resting-State Functional Connectivity in Major Depression: Abnormally Increased Contributions from Subgenual Cingulate Cortex and Thalamus. Biol Psychiat. 62: 429–437.

63. Spielberg JM, Beall EB, Hulvershorn LA, Altinay M, Karne H, Anand A (2016): Resting State Brain Network Disturbances Related to Hypomania and Depression in Medication-Free Bipolar Disorder. Neuropsychopharmacol. 41: 3016.

64. Martino M, Magioncalda P, Huang Z, Conio B, Piaggio N, Duncan NW, et al. (2016): Contrasting variability patterns in the default mode and sensorimotor networks balance in bipolar depression and mania. Proc National Acad Sci. 113: 4824–4829.

65. Cloninger RC, Svrakic DM, Przybeck TR (1993): A Psychobiological Model of Temperament and Character. Arch Gen Psychiat. 50: 975–990.

66. Celikel F, Kose S, Cumurcu B, Erkorkmaz U, Sayar K, Borckardt JJ, Cloninger RC (2009): Cloninger’s temperament and character dimensions of personality in patients with major depressive disorder. Compr Psychiat. 50: 556–561.

67. Öngür D, Farabaugh A, Iosifescu DV, Perlis R, Fava M (2005): Tridimensional Personality Questionnaire Factors in Major Depressive Disorder: Relationship to Anxiety Disorder Comorbidity and Age of Onset. Psychother Psychosom. 74: 173–178.

68. Whitton AE, Treadway MT, Pizzagalli DA (2015): Reward processing dysfunction in major depression, bipolar disorder and schizophrenia. Curr Opin Psychiatr. 28: 7.

69. Woo C-W, Chang LJ, Lindquist MA, Wager TD (2017): Building better biomarkers: brain models in translational neuroimaging. Nat Neurosci. 20: 365–377.

